# Evolutionary Origin of the Mammalian Hematopoietic System Found in a Colonial Chordate

**DOI:** 10.1101/206318

**Authors:** Benyamin Rosental, Mark Kowarsky, Jun Seita, Daniel M. Corey, Katherine J. Ishizuka, Karla J. Palmeri, Shih-Yu Chen, Rahul Sinha, Jennifer Okamoto, Gary Mantalas, Lucia Manni, Tal Raveh, D. Nathaniel Clarke, Aaron M. Newman, Norma F. Neff, Garry P. Nolan, Stephen R. Quake, Irving L. Weissman, Ayelet Voskoboynik

**Affiliations:** Institute for Stem Cell Biology and Regenerative Medicine, and Ludwig Center, Stanford University School of Medicine.; Departments of Biology, Stanford University, Hopkins Marine Station, Pacific Grove, CA 93950, USA.; Department of Physics, Stanford University, CA 94305, USA.; AI based Healthcare and Medical Data Analysis Standardization Unit, Medical Sciences Innovation Hub Program, RIKEN, Tokyo 103-0027, Japan.; Department of Microbiology and Immunology, Stanford University School of Medicine, Stanford, CA 94305, USA.; Departments of Applied Physics and Bioengineering, Stanford University, and Chan Zuckerberg Biohub, San Francisco CA 94158, USA.; Dipartimento di Biologia, Università degli Studi di Padova, Padova, Italy.

## Abstract

Hematopoiesis is an essential process that evolved in multicellular animals. At the heart of this process are hematopoietic stem cells (HSCs), which are multipotent, self-renewing and generate the entire repertoire of blood and immune cells throughout life. Here we studied the hematopoietic system of *Botryllus schlosseri*, a colonial tunicate that has vasculature, circulating blood cells, and interesting characteristics of stem cell biology and immunity. Self-recognition between genetically compatible *B. schlosseri* colonies leads to the formation of natural parabionts with shared circulation, whereas incompatible colonies reject each other. Using flow-cytometry, whole-transcriptome sequencing of defined cell populations, and diverse functional assays, we identified HSCs, progenitors, immune-effector cells, the HSC niche, and demonstrated that self-recognition inhibits cytotoxic reaction. Our study implies that the HSC and myeloid lineages emerged in a common ancestor of tunicates and vertebrates and suggests that hematopoietic bone marrow and the *B. schlosseri* endostyle niche evolved from the same origin.

---

Hematopoietic stem cells (HSCs) are multipotent, self-renewing cells that generate all mature blood and immune cell populations throughout life. In mammals, the blood-forming organ is the bone marrow, where HSCs reside in specialized niches that support their self-renewal and maintenance of an undifferentiated state. Since the identification of HSCs by prospective isolation^1^, several model systems have been studied to elucidate HSC biology^2–8^. While there are comprehensive studies on vertebrate HSC self-renewal, differentiation, physiological regulation and niche occupation, relatively little is known about their evolutionary origin and niches. To gain insight into the evolutionary origin of the mammalian hematopoietic and immune system, and identify fundamental cell and molecular mechanisms underlying HSCs homeostasis and regeneration we characterized the blood and immune system of the colonial tunicate *Botryllus schlosseri*.

Tunicates are marine invertebrates in the phylum Chordata that share the chordate characteristics of a notochord, dorsal neural tube, and gill slits in their free-swimming larval stage, but lose most of these features during metamorphosis into sessile adults^9,10^. In *B. schlosseri*, after metamorphosis, a colony is formed via asexual reproduction consisting of multiple individuals, called zooids, through a stem-cell-mediated budding process that continues throughout life11. Every week, the developed buds replace the parent zooids that undergo synchronized programmed cell death^12^ (Movie S1). Each mature zooid has digestive and respiratory systems, a tube-like heart, siphons, an endostyle, a neural complex, an ovary and testis (Fig. 1a). A vasculature with circulating blood cells connects all zooids and buds within the colony (Fig. 1b).

*B. schlosseri* undergoes a natural transplantation process with genetically compatible colonies, a feature unique among chordates. When two colonies touch, they either fuse their vasculature and share blood, or reject^13,14^ (Fig. 1c; Movie S2). This self-nonself recognition process is controlled by *BHF*, a single polymorphic histocompatibility gene15. Fusion requires at least one shared *BHF* allele: upon fusion, circulating germline and somatic stem cells from both colonies compete for dominance and contribute to the formation of new buds^11,15–18^. Additionally, the budding cycle in one of the chimeric partners often fails to develop^19^. This developmental failure is an immune-cell based rejection of bud cells that operates within a *BHF* histocompatible chimera. It involves inflammation and recruitment of cytotoxic cells, and is thus comparable to mammalian chronic rejections^19^.

**Figure 1.**
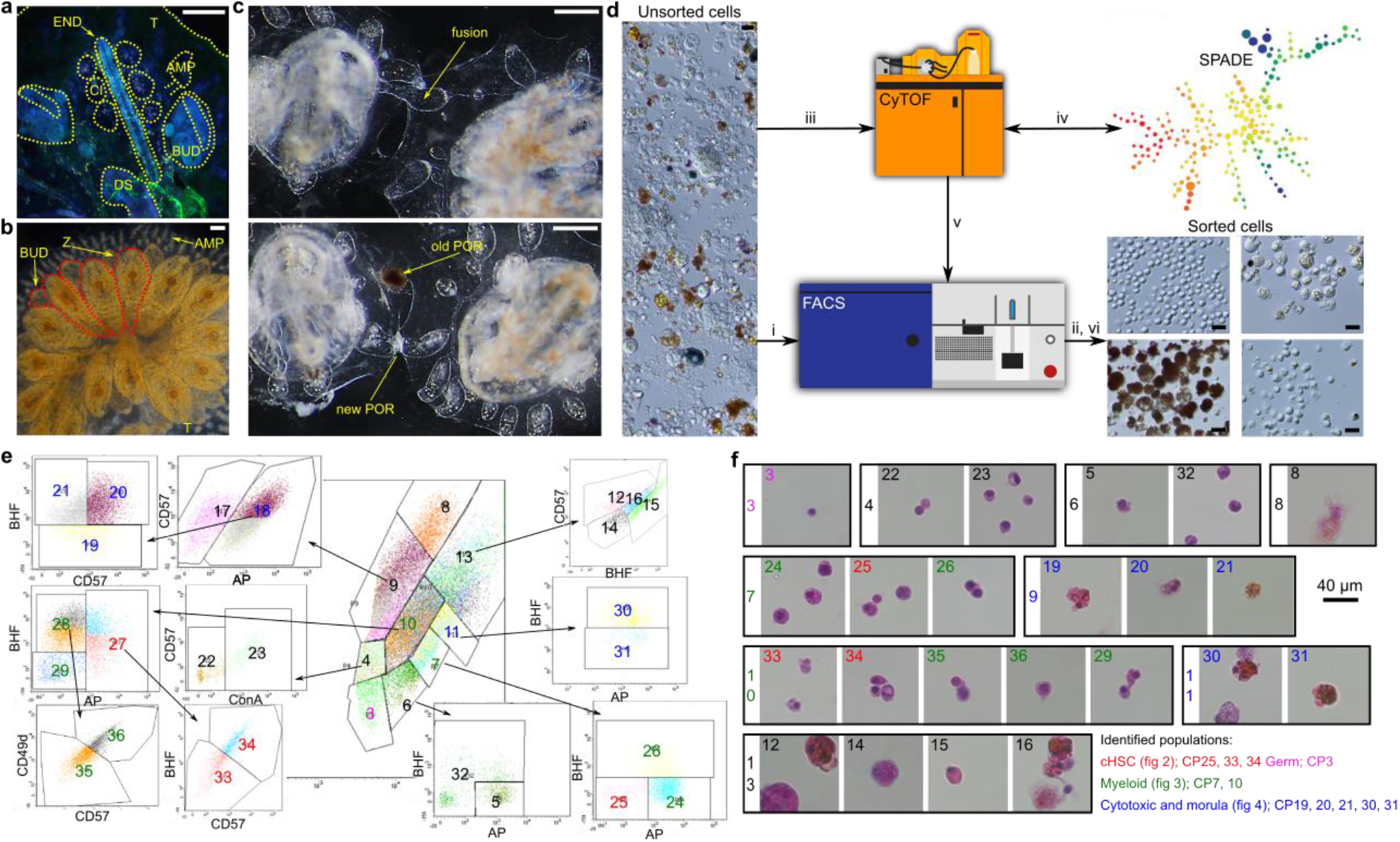
*B. schlosseri* Anatomy and Cell Sorting Workflow. **a,** Fluorescent image of a zooid (dorsal view), DAPI labeled and acquired with auto-fluorescence. The primary bud (BUD), vasculature (V) connecting to an ampulla (AMP), endostyle (END), cell islands (CI), digestive system (DS) and tunic (T) are marked. **b,** Developing buds (BUD) in a colony and the older generation parental zooids (Z), connected to each other by vasculature which terminates with ampullae (AMP). **c,** Live imaging of colonies undergoing fusion (top) and rejection (bottom), arrows point to fused vasculature and points of rejection (POR). Scale bar 0.2 mm. **d,** Outline of cell purification process. Unsorted cells (light microscopy) are loaded into a FACS (i) and sorted resulting in morphological observation (ii; Fig. S1). Cells were labeled by diverse markers and screened by CyTOF (iii). Based on SPADE cluster analysis (iv) markers were selected for FACS gating (v) before a final sort was performed (vi; **e**). (**e**) Sorting panels of 34 cell populations. Central panel is FCS/SSC from which additional populations were differentiated. **f,** H&E staining of the end point cell populations isolated in a rectangle of original population by FCS/SSC. Color key (reflected in subparts **e-f**) for different populations identified in this study and the figures that further describe them.

Charles Darwin suggested that tunicates may be the key to understanding the evolution of vertebrates, and recent genomic data has revealed that they are indeed the closest sister group of vertebrates^10,20^. Furthermore, comparing the *B. schlosseri* genome with those of several vertebrates, revealed that *B. schlosseri* has numerous hematopoietic system related genes, many shared with mammals^21^.

To characterize the *B. schlosseri* hematopoietic and immune system at the cellular and molecular levels, we deployed advanced genomics, live imaging and classical immunological methods developed specifically for the study of the mammalian immune system including multi-parameter flow cytometric isolation of cellular populations, cell sorting, transplantation assays as well as phagocytosis and cytotoxicity measurement.

## Prospective isolation of candidate HSCs and myeloid lineage

To understand the composition of the *B. schlosseri* hematopoietic and immune systems, we separated and analyzed its cell populations (Fig. 1d-f). Using fluorescence-activated cell sorting^22^ (FACS) we first isolated 11 cell populations based on their size (FSC), granularity (SSC) and natural auto-fluorescence (Fig. S1). We then screened a large assortment of antibodies and reagents by CyTOF and FACS, to identify markers that show differential binding to distinct cell populations (Tables S1, S2; Fig. S2). Based on cluster analysis, markers were selected for FACS gating (Fig. 1d). Using CD49d, CD57, Concanavalin-A, BHF (a mouse serum that was developed against the BHF protein), alkaline phosphatase (AP), together with size, granularity, and natural auto-fluorescence, we succeeded in defining a total of 34 cell populations - 24 of these are the hierarchical endpoint populations of our gating strategy (Fig. 1e-f, S3).

To characterize the *B. schlosseri* cell populations (CP) isolated by FACS, we sequenced the transcriptome of the 24 sorted cell populations (Fig. 1e-f, S4, Table S3). Many of the populations have visually similar gene expression patterns (for example: CP3, 5 and 23; and CP25, 33 and 34), and in general, samples that were adjacent in FACS-space are also similar in expression-space (Fig. S4, over 67% of neighbors in agreement) indicating a correlation between gene expression profile, morphology and marker expression. The cluster of cell populations CP25, 33 and 34, had 235 differentially upregulated genes known to be expressed in the vertebrate blood and hematopoietic systems^23^ (Figure S3). The Geneset Activity Analysis of the Gene Expression Commons^24^ was developed and applied to this gene set, to allow simultaneous comparison of multiple genes to expression data from 39 distinct and highly purified mouse hematopoietic stem, progenitor and differentiated cells. This analysis revealed significant expression overlap with mammalian hematopoietic stem and progenitor cells and with cells of myeloid lineages (Fig. 2a, Table S4).

**Figure 2.**
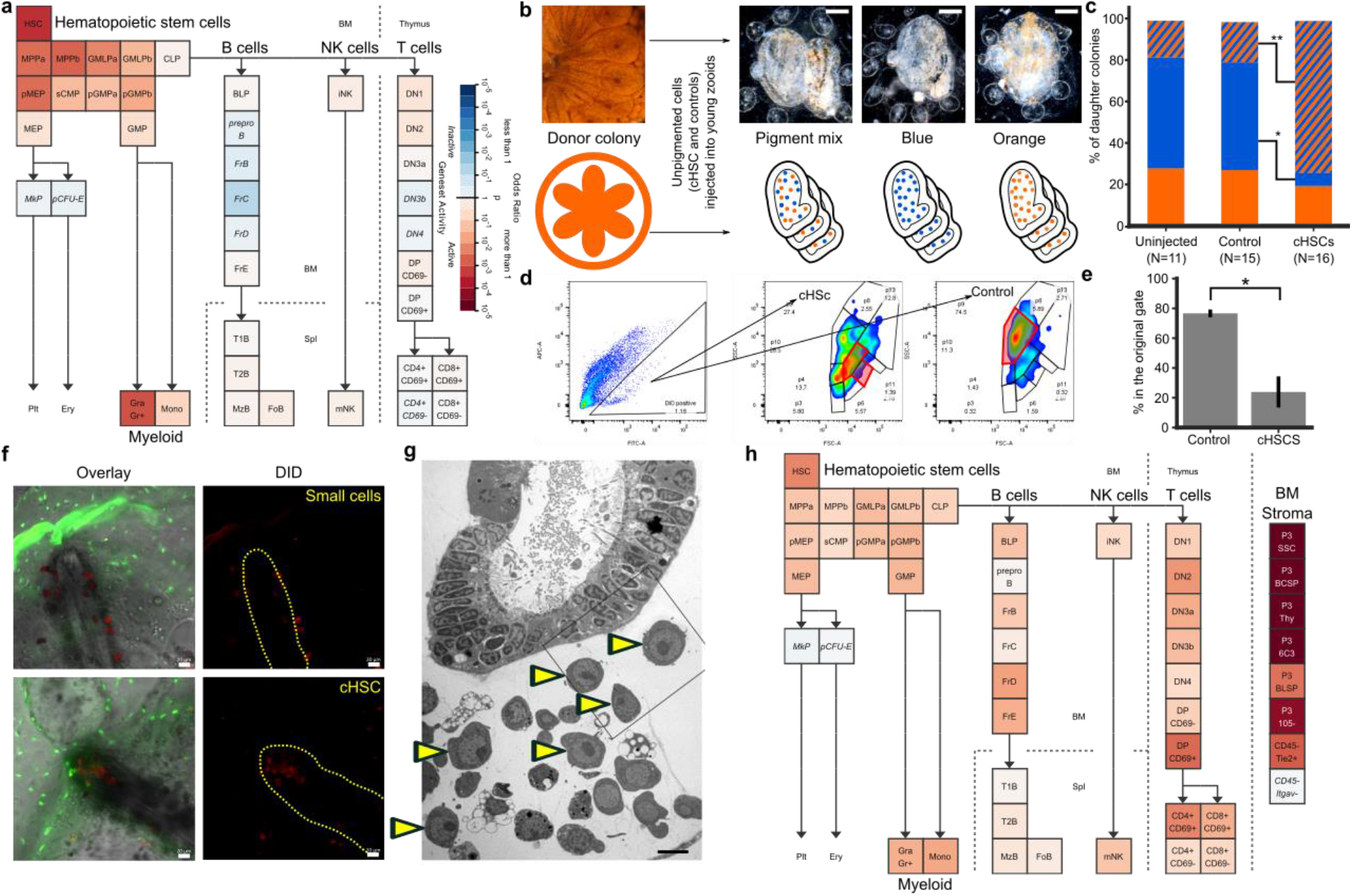
Multilineage Differentiation Capacity, Homing Sites of HSC and their niche. **a,** Geneset Activity Analysis of 235 genes upregulated in CP33, 34 and 25 using the Gene Expression Commons tool on a mouse hematopoiesis model. The enriched populations are HSCs and the myeloid lineage. **b,** Candidate HSCs (cHSC) and a control cell population from an orange pigmented donor colony were transplanted into compatible recipient colonies with blue, orange, or mixture of the two pigmented cells. Upper panel shows light microscopy, lower panel is an illustration. Scale bar 0.2 mm. **c,** Significant reduction of the ratio of blue colonies and significant upregulation of the ratio of mixed pigmented colonies (Fisher’s exact test, * P<0.05, ** P<0.01), 20 days post-cHSC transplantation. **d-e,** cHSC and a control cell population were labeled with DiO and transplanted into compatible colonies. Three weeks after transplantation DiO+ cells from the recipient colonies were analyzed by FACS (left). The majority of the transplanted cells from the control group were detected in the original gate (in red). Only 24% of the cHSC transplanted cells were detected in their original gate, the rest were detected in diverse cell populations. Error bars - sd. **f,** cHSC population and a control population were isolated, labeled with DiD and transplanted into CFSE labeled compatible colonies, five to ten days afterwards, only the cHSC populations migrated and aggregated in the endo-niche. Scale bar 20 μm. **g,** Electron microscopy section of the endostyle and sub-endostylar sinus (endo-niche). Yellow arrows indicate cells with hemoblast (HSC) morphology. Scale bar 5 μm. **h,** Geneset Activity Analysis by Gene Expression Commons reveals significant similarities between the endostyle gene expression and the mouse bone marrow stromal cells (scale same as in **a**).

To measure the ability of the candidate HSC (cHSC) population and the control cell populations to differentiate to other cell types, we developed two transplantation assays: i) a pigment cell based differentiation assay (donor chimerism) (Fig. 2b-c), and ii) a FACS analysis based differentiation assay (Fig. 2d-e). The colors of the pigment cells in *B. schlosseri* are genetically determined^25,26^. Pigment cells circulate in the colony vasculature, are identified by FACS as CP8, and are linked to the hematopoietic lineages by gene expression (Fig. S5b).

Cells were transplanted from orange *B. schlosseri* donor colonies, to compatible recipients with three different pigmentation patterns: orange colonies, blue colonies, or mixed blue-orange pigmented colonies (Fig. 2b). Candidate HSCs were sorted by FACS, and the cell population closest to the pigment cells based on FACS and the molecular analysis (CP18; Fig. 1e-f, S4), served as the control. Twenty days following transplantation the recipient colonies transplanted with the control (CP18) showed no significant difference in the ratio of orange, blue and mixed pigment colonies compared to uninjected colonies (Fig. 2c). However, the recipient colonies transplanted with the cHSC population had a significant reduction in pure blue colonies (53% to 6%) and an upregulation of the mixed population colonies (20% to 75%). This demonstrates that the unpigmented cHSC population from the orange donor colony differentiated into orange pigment cells within the recipient colonies, creating donor chimerism.

Next, we used FACS analyses to assess the differentiation potential of transplanted cells. In this set of experiments the cHSC population and control population (CP18) were isolated from donor colonies, labeled with a fluorescent membrane dye (DiO), and transplanted into compatible recipient colonies. Three weeks following transplantation DiO+ cells from the recipient colonies were analyzed by FACS. As demonstrated in Fig. 2d (right), more than 70% of the transplanted cells from the control group were detected in their original FACS gate. In contrast, only 24% of the cHSC transplanted cells, were detected in their original gate, the rest of the labeled cells were detected in diverse gates of distinct *B. schlosseri* cell populations (Fig. 2d-e). This demonstrates the multilineage differentiation capacity of the cHSC population.

## The bone marrow and the endostyle share gene expression and function

To identify cHSC niches we transplanted labeled cell populations and traced them *in-vivo* utilizing the transparent body of *B. schlosseri*. The cHSC population and a control population (CP3) were isolated, labeled with a lipophilic dye (DID) and injected into labeled (CFSE) compatible colonies (Fig. 2f). Following transplantation, DID-labeled cHSC populations migrated in the recipient colony and aggregated in the two known *B. schlosseri* stem cell niches; the endostyle niche which is located at the anterior sub-endostylar sinus (endo-niche), previously identified as a somatic stem cell niche, and the cell islands niche, considered a germline stem cell niche^18,27^ (Fig. 2f, S6a). Moreover, blood cells with hemoblast (HSC) morphology are abundant in the endo-niche (Fig. 2g, S6b). This experiment demonstrated that the cHSCs localize to known stem cell niches, specifically to the endo-niche. Transplanted cells from the control population (CP3) did not home to the endostyle stem cell niche, but were found in the cell islands niche (Fig. S6a). Interestingly, the control population, which does not express a gene signature of the hematopoietic system, did have a germline gene expression signature and localized to the known germline stem cell niche (Fig. S5c, S6a).

To further characterize the cHSC niche, we generated comprehensive transcriptome sequence data from ten samples of dissected endostyles and compared them to the transcriptome of 34 whole colonies. Genes that were significantly upregulated in the endostyle with homology to human or mouse genes were analyzed by GeneAnalytics^23^, revealing a shared expression of 327 genes between the endostyle and human hematopoietic bone marrow (Fig. S6c, Table S4). This finding was further supported by Gene Expression Commons, which revealed significant similarities in gene expression patterns between the endostyle and mouse bone marrow stromal cells^5,24^ (Fig. 2h). The similar genetic signature between the *B. schlosseri* endostyle and the hematopoietic bone marrow and the high abundance and close proximity of cHSCs within the endo-niche (Fig. 2g; S6b), suggest that the endostyle is the hematopoietic organ.

## Discovery of a myeloid lineage phagocytic population

To identify the *B. schlosseri* myeloid lineage phagocytic cell populations, we used three different *ex-vivo* phagocytosis assays: i) phagocytosis of fluorescent beads, ii) phagocytosis of a fluorescently labeled marine bacteria; *Vibrio diazotrophicus*, and iii) allogeneic phagocytosis, where we tested the capability of cells from different colonies to engulf allogeneic cells. FACS was used to identify the phagocytic cells in each of the *ex-vivo* assays and to track its cell population (Fig. 3a). Two previously described populations of phagocytic cells were identified: the amoebocytes (within CP4 and 18) and the large phagocytes28 (within CP13), and also a third population, the candidate myeloid cell population (within CP7 and 10) (Fig. 3a, S7a). We then isolated the three main phagocytic populations and a small cell population (CP3) as a control, and incubated each one with fluorescent beads to validate engulfment capacity of each population. While only 1% of the control cells engulfed the beads, 7% of the large phagocytes and 20% of both the amoebocytes and myeloid cells engulfed the beads (Fig. 3b-c). Confocal microscopy and ImageStream analysis confirmed that the three phagocytic populations assayed engulfed the beads (Fig. 3d, S7b). The myeloid cells were found to be the main contributors to phagocytosis, as evidenced by >40% contribution to each of the phagocytosis assays (Fig. 3a). It appears that the large phagocytes contribute mainly to allogeneic phagocytosis, compared to other assays (Fig. 3a). These *ex-vivo* functional phagocytosis assays revealed three major phagocytic populations in *B. schlosseri*.

**Figure 3.**
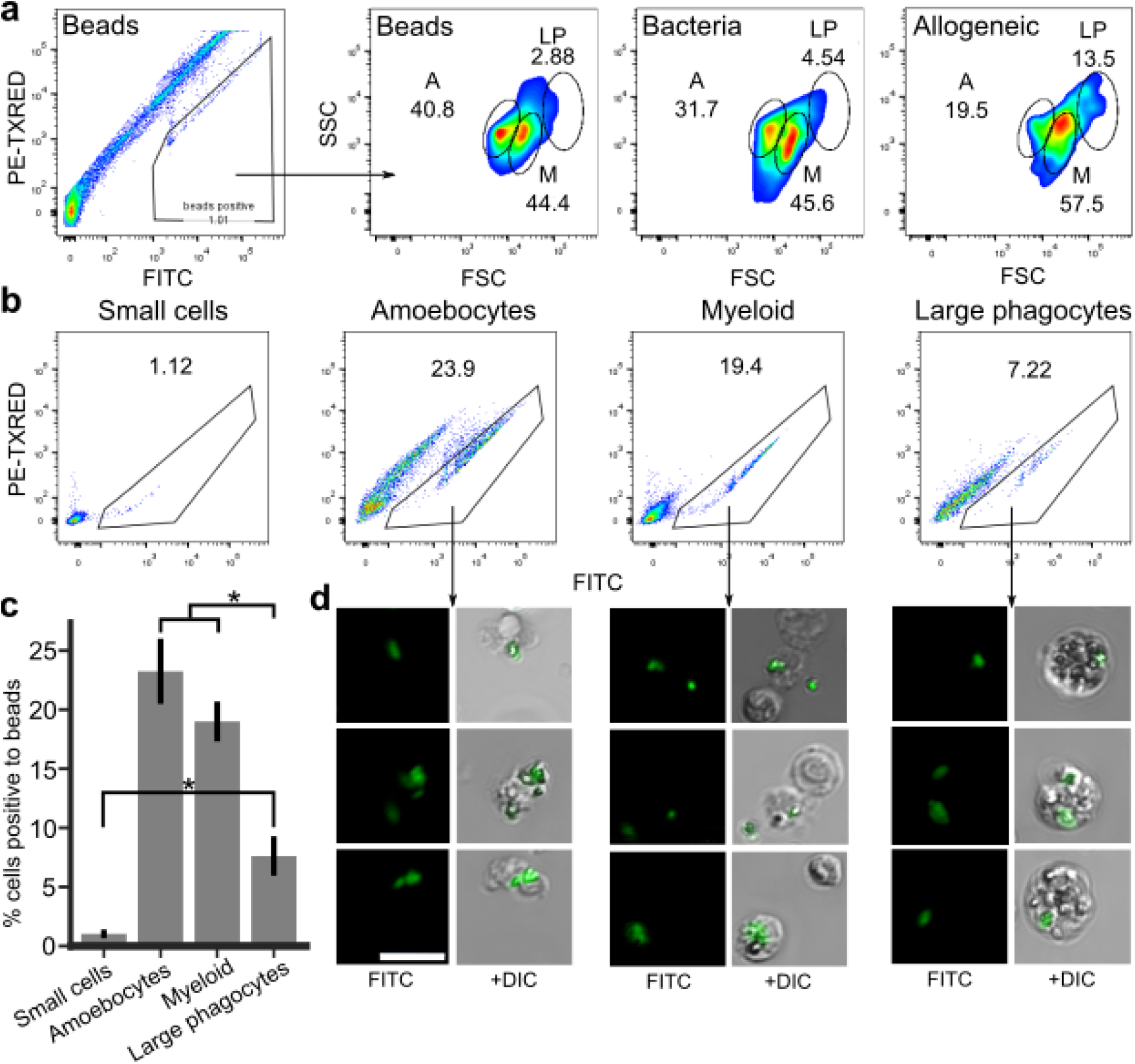
Discovery of a Myeloid Lineage Phagocytic Population. **a,** FACS analysis of *B. schlosseri* cells that are fluorescently positive in one of three phagocytosis assays performed: (first and second) phagocytosis of green fluorescent beads, (third) phagocytosis of *Vibrio diazotrophicus* labeled with AF647, and (fourth) allogeneic phagocytosis. Three phagocytic populations identified: amoebocytes (A), myeloid cells (M), and large phagocytes (LP). **b,** FACS analysis of green fluorescent beads phagocytosis of sorted populations. **c,** Amoebocytes, myeloid cells, and large phagocytes all had significantly higher engulfment rates than the small cell population. In addition, amoebocytes, myeloid cells had significantly higher cell percentages than the large phagocyte population (ANOVA, * P<0.05). **d,** Representative images of the three phagocytic populations after engulfment of beads. Scale bar 20 μm.

## BHF-mediated self-recognition inhibits cytotoxicity

In mammals, cellular immune responses are mediated by phagocytosis: the engulfment of target cells, and cytotoxicity: the direct killing of the target cells. To test whether a cellular recognition response can take place *in-vivo* during rejection, whole colonies were differentially labeled, set near each other and monitored by live imaging (Fig. 4a, Movie S3). Direct contact between allogeneic cells was detected during rejection within the points of rejection (POR; Fig. 4a, lower right). This observation demonstrates that allogeneic cellular recognition can be facilitated *in-vivo* in the process of colony rejection.

**Figure 4.**
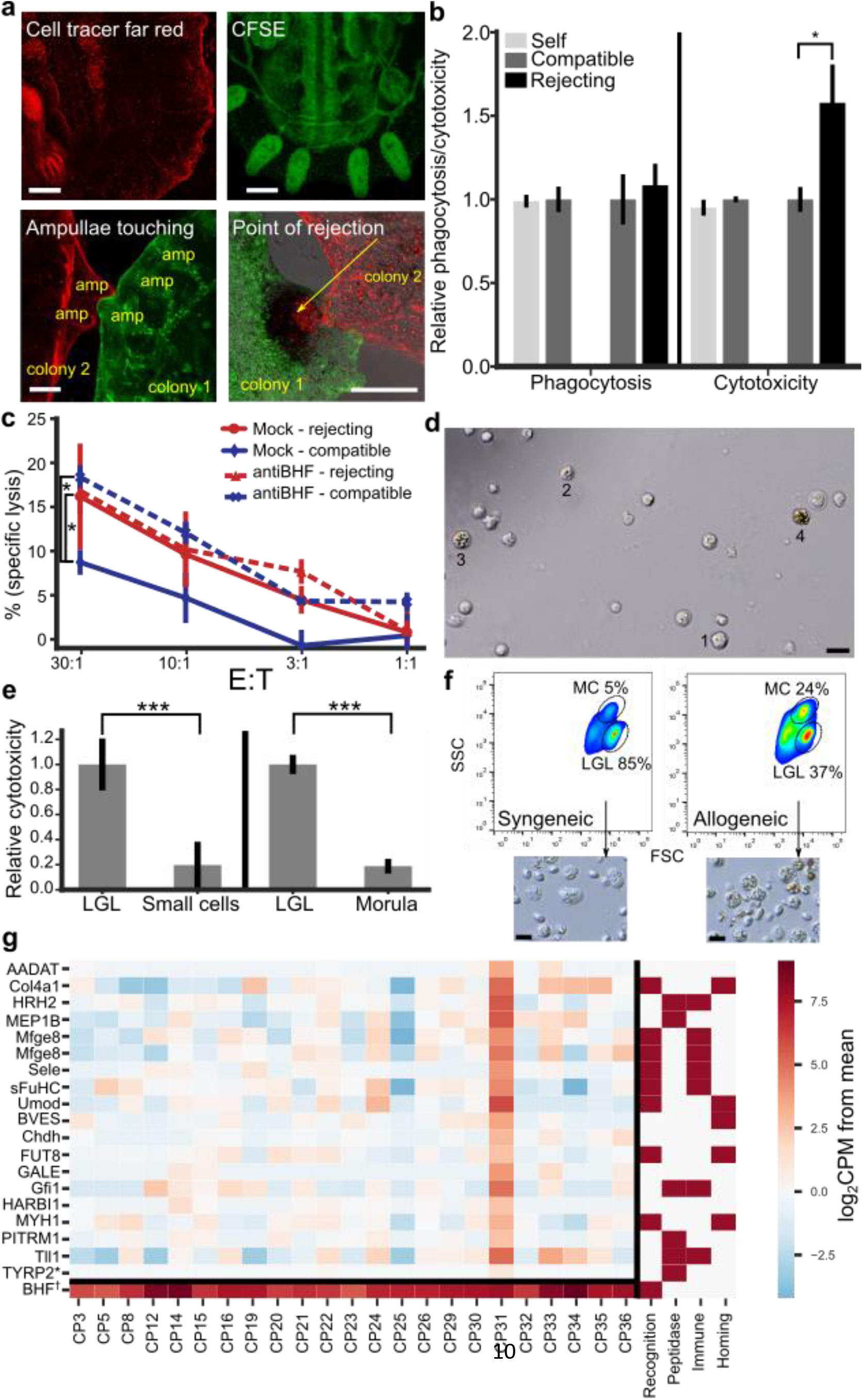
Allorecognition is mediated by cytotoxic cells through BHF recognition. **a,** Colony labeled with cell tracer far red (top left), and with CFSE in green (top right). Ampullae touching (bottom left), and in points of rejection (bottom right) the mixture of the two colors of cells can be observed in the area of the necrotic tissue (dark area, yellow arrow). Scale bar 0.2mm. **b,** Phagocytosis and cytotoxicity assays were set between compatible, incompatible, or self. Significantly upregulation of cytotoxicity between cells taken from incompatible colonies are observed. **c,** Blocking of BHF with anti BHF serum. Compatible colony target cells in blue and incompatible colony target cells in red. Mock serum-solid line and anti-BHF serum-dashed line. The blocking of BHF significantly upregulated the cytotoxicity of compatible targets compared to mock serum. **d,** Observation of Isolated large granular lymphocyte-like (LGL) cells after two days *ex-vivo*. 1-morphology of original isolated cells, 2-3-granular and light pigmented cells, and 4 - granular pigmented morula cell (MC). **e,** Cytotoxicity assays of isolated LGL cells compared to small cells, and to isolated MC. In both cases the LGL cells had significantly higher cytotoxicity compared to the other cell types. **f,** LGL cells were isolated and incubated overnight either in syngeneic or in allogeneic challenge. FACS analysis of LGL and MC distribution and caption of light microscopy images of cells following incubation. Scale bar 20 μm. **g,** Representation of homolog annotated genes differentially upregulated by LGL enriched population CP31. *TYRP2 (homolog of Phenoloxidase) is not differentially expressed, but is 7-fold higher in CP31. †BHF expression levels in the cell population are included (in log counts per million). Color scaling counts per million in log2, red high, blue low. Right side is a table indicating gene association with immunity functions. Standard Deviation; ANOVA *P<0.05, ***P<0.005.

Next, we performed allogeneic phagocytosis and cytotoxicity assays. Assays were set between cells taken from compatible colonies, incompatible colonies (rejecting), or self (Fig. 4b). Confocal microscopy was used to validate phagocytosis and for cytotoxicity, specific lysis was validated by increasing ratios between effector to target cells (Fig. 4c, S8a-b). Similar levels of phagocytosis were detected between cells that were isolated from the same colony (self), compatible colonies or incompatible colonies. A significant increase of cellular cytotoxicity was detected between cells that were derived from incompatible colonies, compared to cells from the same colony or compatible colonies (ANOVA P<0.05; Fig. 4b, S8b). These results demonstrate that cytotoxicity is the immune effector mechanism of the cellular allogeneic response.

In a third set of experiments we tested whether blocking of BHF would affect the cytotoxic reaction. Cytotoxicity assays between compatible and incompatible colonies revealed significantly higher cytotoxicity between incompatible colonies with mock serum (Fig. 4c). Blocking of BHF with serum containing polyclonal anti-BHF antibodies that bind live *B. schlosseri* cells (Fig. S3, Table S2), enhanced cell lysis between compatible colonies to the level observed in incompatible colonies (Fig. 4c). This treatment did not affect the level of cell lysis observed in incompatible colonies (Fig. 4c). These results revealed that: (i) BHF is a major histocompatibility factor essential for self-recognition; (ii) BHF recognition prevents cell lysis by allogeneic cytotoxic cells via inhibition of cellular cytotoxicity.

## Cytotoxic large granular lymphocyte like cells

We did not identify a cell population with a clear mammalian cytotoxic gene expression signature. Morula cells (MC), cells that contain phenoloxidase, accumulate in rejection points and have been proposed as cytotoxic cells that mediate rejection^28–30^. We detected MC in population CP18 (mother population of CP19, 20 and 21; Fig. 1e-f, S3). However, the number of MC in these populations was lower than expected, prompting us to look for a candidate precursor cell. We studied the large granular lymphocyte-like (LGL) cells (enriched in CP31) as a potential candidate, since its morphology resembles Natural Killer (NK) cells, the innate immunity cytotoxic cells of mammals, which originally were characterized as large granular lymphocytes^31^. Indeed, *ex-vitro* experiments revealed a transition of purified LGL cells into MC after two days *in-vitro* (Fig. 4d, labeled 1-4). Based on these observations we hypothesized that the LGL cells are the cytotoxic MC which become pigmented granular MC following activation.

To test this hypothesis, we performed cytotoxicity assays that compared the ability of MC and LGL cells to induce cytotoxicity. This set of experiments revealed that isolated LGL cells had significantly higher cytotoxicity compared to isolated MC and to small cells (CP3; Fig. 4e). We then tested whether LGL cells become MC upon activation. LGL cells were isolated, incubated overnight in either a syngeneic (self) or an allogeneic challenge. While, the majority of the LGL cells that were incubated with syngeneic cells remained in the LGL population (>60%), less than 10% became MC (Fig. 4f, S8c). On the other hand, less than 40% of the LGL cells that were incubated with allogeneic cells remained in the LGL population, the number of cells in the MC population almost tripled (Fig. 4f, S8c). The transition from LGL morphology to MC was further validated by light microscopy (Fig. 4f lower). Upon activation, LGL change their morphology, develop granularity and pigmentation, presumably due to phenoloxidase activation, and become MC (Fig 4f, S8d). This set of experiments demonstrates that the LGL cells are the cytotoxic cell population. The term cytotoxic morula cell will be used to describe the LGL cells.

Analysis of the genes differentially upregulated by the highly enriched cytotoxic morula cells population (CP31), revealed 85% tunicate specific genes with no human or mouse homologs. Interestingly, among the 18 differentially upregulated genes that do have sequence homology to genes in vertebrates, 14 are associated with at least one of the following functions: cellular recognition, cytotoxicity or peptidase activity, leukocyte homing and general immune response (Fig. 4g). These genes are involved in toxin transport, leukocyte cellular adhesion, and heterothallic cellular adhesion through membrane proteins - all processes characteristic of cytotoxic cells. CP31 also expresses tyrosinase, the vertebrate homolog gene to phenoloxidase which is one of the main enzymes found in MC^29,32^, 7-fold higher compared to the other cells. While we did not find a *B. schlosseri* cell population that had a significant lymphoid lineage signature, we did find that the MC enriched CP19 has expression resemblance to mouse T-cells, B cells, and immature NK cells through Geneset Activity Analysis by Gene Expression Commons (Fig. S5d), this list doesn’t include any gene that is associated with known adaptive immunity function.

## Evolution of hematopoietic and cellular immune systems

In mammals and reptiles the bone marrow is the main center of hematopoiesis, but the site of hematopoiesis in other species reveals a list of diverse organs and tissues ^8,33–39^. In *B. schlosseri*, we identified a somatic stem cell niche in the anterior subendostylar sinus^18^. Here we discovered that the molecular signature of the endostyle is similar to mammalian hematopoietic bone marrow and that the specific somatic stem cells that reside in and home to this niche are HSCs. Additional cellular and molecular comparison of diverse hematopoietic niches may reveal conserved elements that were maintained throughout evolution, essential for hematopoiesis.

Among the genes expressed in *B. schlosseri* isolated cell populations, we found high enrichment for gene sets predominantly expressed in human HSC and myeloid populations, consistent with previous studies^21,40–45^. In mammals, the Major Histocompatibility Complex (MHC) is the tissue-antigen that allows the immune system to recognize and tolerate ‘self’. Histocompatibility is governed by cytotoxic T-cells and NK cells. NK cell cytotoxicity is inhibited when Killer Inhibitory Receptors (KIRs) recognize self-MHC, and is activated when cells do not express self-MHC; ‘missing-self’^46^. Immune functional assays further revealed that cellular rejection between genetically incompatible colonies is mediated by cytotoxicity and that BHF is a self-recognition inhibitory factor for cytotoxic cells. Similar to NK inhibition by MHC, blocking of BHF revealed that the self-BHF recognition is a major inhibitory mechanism of cytotoxicity. These results and the observation that colonies sharing at least one *BHF* allele fuse^15^, demonstrate that like in NK recognition, the cellular cytotoxicity mechanism in *B. schlosseri* is based on ‘missing self’. We hypothesize that recognition of self BHF by inhibitory receptors expressed on the *B. schlosseri* effector cells, is the mechanism of self-recognition. This suggests an education mechanism comparable to the KIR repertoire for self-MHC^47^, during maturation of the *Botryllus* cytotoxic cells.

The *B. schlosseri* hematopoietic and immune systems combine vertebrate and invertebrate features (Fig. 5). *B. schlosseri* shares with vertebrates a myeloid lineage including cells that take part in phagocytosis. It also has amoebocytes and large phagocytes with morphologies resembling invertebrate cell types^48,49^. The *B. schlosseri* cytotoxic morula cells carry imprints reminiscent of vertebrates’ lymphocytes but mainly express tunicate-specific gene repertoire (Fig. 4g, S5d). Several studies describe morula-like cells with phenoloxidase activity in other invertebrate species^32,49^. Studying novel gene sets that the cytotoxic morula cells express, will most likely reveal novel mechanisms to target pathogens and delimit self from non-self.

**Figure 5.**
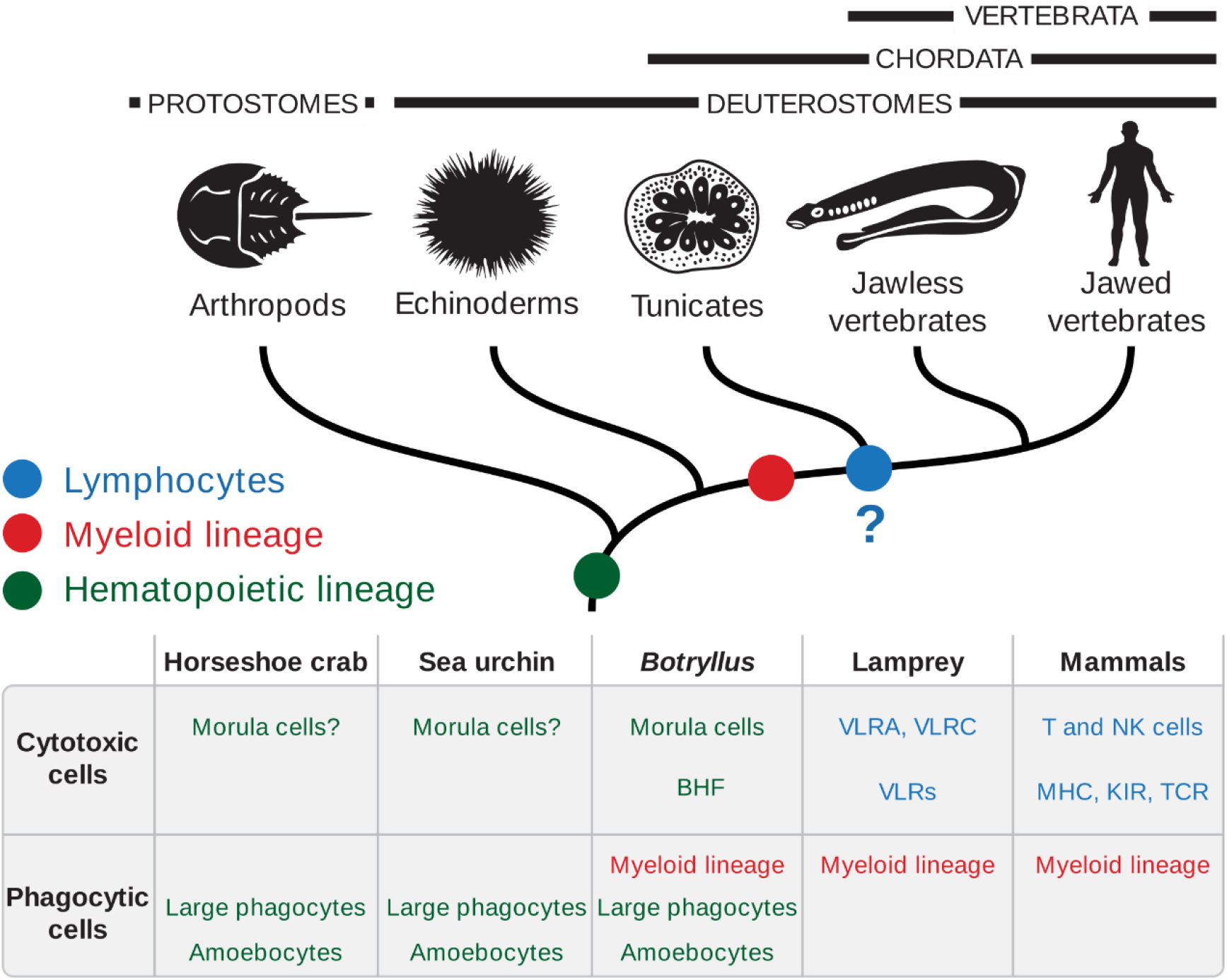
Evolution of cellular immune system. Proposed evolution of the cellular immune system, mainly cytotoxic and phagocytic cell lineages. Myeloid lineage evolved before the branching of the vertebrata from tunicates, in red. Whether the lymphoid lineage evolved in the common ancestor of tunicates and vertebrates was not deciphered in the current research (in blue); there is a cellular population and additional molecular analysis suggesting lymphoid cells, mainly undifferentiated. In cytotoxic cells second line are the cellular recognition molecules: Immunoglobulin superfamily MHC, KIRs and TCR, leucine rich repeats receptors of the VLRs, and the BHF.? – Represent missing functional validation.

Our paper describes a thorough cellular, molecular, and functional characterization of the *B. schlosseri* hematopoietic and immune systems, including the identification of the HSCs and progenitors, myeloid cell lineage within three phagocytic cell populations, cytotoxic cells that induce cellular tissue rejection, and finally the discovery of the hematopoietic organ showing that it is similar to human hematopoietic bone marrow. Adapting the tools and methods developed here to diverse organisms will enable the study of immune systems diversity and will reveal novel defense mechanisms in nature.

## Acknowledgments

The authors would like to thank Chris Lowe, Ivan Dimov, Seth Karten, Chris Patton, Judy Thompson, Patty Lovelace, Ronnie Voskoboynik, Nathaniel Fernhoff, Pauline Chu, Jonathan Tsai, Kipp Weiskopf, Matan Oren, John Lee, Barbara Compton, Kevin Uhlinger, Tejaswitha Naik, and Terry Storm for invaluable technical advice and help. This study was supported by NIH grants R56AI089968, R01AG037968 and RO1GM100315 (to I.L.W., S.R.Q., and A.V.). The Virginia and D. K. Ludwig Fund for Cancer Research, a grant from the Siebel Stem Cell Institute and Steinhart-Reed grant (to I.L.W.). L.M. was supported by PRIN - Prot. 2015NSFHXF. The work of B.R. was supported by the Postdoctoral Fellowship of the Human Frontier Science Program Organization, the NIH Immunology training grant 5T32AI07290-28 and the NIH Hematology training grant T32 HL120824-03.

## Author contributions

Conception and design B.R., A.V., M.K., S.R.Q., I.L.W.; Mariculture K.J.I, K.J.P.; Flow cytometry and sorting B.R.; CyTOF screening and cluster analysis B.R., S.Y.C., G.P.N.; RNA isolation and library preparation B.R., K.J.P., R.S., A.V.; Sequencing J.O., G.M., N.F.N.; Sequencing analysis and developed analytical tools M.K., J.S, A.M.N., S.R.Q.; Immunological assays B.R.; Microscopy and experimental design B.R., D.M.C., K.J.I., D.N.C., A.V; Electron microscopy L.M.; Transplantation B.R.; Writing of manuscript B.R., A.V., M.K., T.R., K.J.P., I.L.W.; Technical support and conceptual advice N.F.N., D.M.C., A.M.N., T.R., G.P.N., S.R.Q., A.V., I.L.W.

## The authors declare no conflict of interest

**Accession number for the sequencing data is PRJNA414486**

## Methods

### Animals

Mariculture procedures have been described previously^50^. Briefly, *Botryllus schlosseri* colonies were collected from the marina in Monterey, California. Individual colonies were tied to 3x5-cm glass slides and placed 5 cm opposite another glass slide in a slide rack. The slide rack was placed into an aquarium, and within a few days the tadpoles hatched, swam to the settlement slide, and metamorphosed into the adult body plan (oozooid). Single oozoids are then transferred to individual slides to provide colonies for lab experiments.

### FACS analysis

For cell isolation, colonies were dissociated with a fine blade into cell suspensions and filtered through a 40 μm mesh using a sterile 1 ml syringe pump, washed and collected in staining media: 3.3x PBS, 2% FCS and 10 mM Hepes. After gating on Propidium Iodide (PI), negative cells (using two dimensional plots due to natural fluorescence of *B. schlosseri* cells), were analyzed using size - forward scatter (FSC) and granularity - side scatter (SSC) panel on log scale using BD FACS Aria-II. Additional gating analysis of the 11 population was then done using the natural autofluorescence of the cells (Fig. S1). These purified populations were analyzed according to morphology by light microscopy and defined according to previous cellular work characterizing *B. schlosseri* cell populations^14,28^. 34 gated populations were analyzed following cell labeling. 5 million cells were suspended in 200μl of staining media prior to FACS analysis: Alkaline Phosphatase (AP) Live Stain (Life Technologies A14353) 1μl, CD49d PE-Cy7 (BioLegend 304313) 1μg, CD57 Pacific Blue (BioLegend 322316) 0.25μg, Concanavalin-A (ConA) AlexaFluor-633 (Sigma) 2μg, mouse anti-BHF serum 1:100 and anti-mouse secondary Cy5-Cy7 (SantaCruz). The specific excitation laser and optical filter for emission measurements was as follows [excitation laser in nm and filters stated as long pass (LP) and band pass (BP)]: 488 nm(505LP 530/30BP)-AP, 488 nm(755LP 780/60BP)-CD49d, 405nm(450/50BP)-CD57, 633 nm(660/20BP)-ConA, 633 nm(755LP 780/60BP)-BHF. Cells were sorted into 18 well flat μ-Slides coated with poly-L-Lysine (ibidi) for live imaging. CP17 was composed of dead cells and cell debris, and was therefore excluded.

### Colony labeling and microscopy

Juvenile colonies were labeled with distinct fluorescent stains (CellTrace CFSE Green, Invitrogen #C34554; CellTrace FarRed, Invitrogen #C34572) and aligned, (or positioned) in controlled fusion/rejection reactions. For allorecognition assays, colonies were labeled using CFSE dye (5 mM stock solution) and Far Red dye (1 mM stock solution) in a dilution of 1 μl of dye to 1 ml of filtered seawater. Naive oozooids were then bathed in this solution for 60 minutes, allowed to recover then washed and placed in μ-Dishes (ibidi μ-Dish 50mm-low uncoated) for observation. Images were obtained using a confocal fluorescence microscope (LSM700 Axio Observer.Z1, ZEISS).

For Hematoxylin and Eosin (H&E) staining: cells were incubated overnight on glass slides coated with Poly-L-lysine (Sigma) and fixed by 4% PFA in 0.1M MOPS in 0.5M NaCl, pH 7.5 for 10min and washed with PBS. The slides were stained with Harris Hematoxylin for 5 minutes and 2% Eosin Y for 1 minute.

### CyTOF mass cytometry screening

Live cells were labeled for 30 minutes on ice in staining media (3.3x PBS, 2% FCS and 10 mM Hepes) with different antibodies labeled with elemental isotopes (Table S1), followed by two washes with staining media. After washing cells were fixed in 4% PFA in 1x PBS, washed once with staining media and then incubated at room temperature for 20 min in an iridium-containing DNA intercalator (Fluidigm) in 1.5% paraformaldehyde in 1x PBS. Prior to measurement on a mass cytometer, cell samples were washed once with staining media and twice with water^51^ (Fluidigm). Analysis of the FCS files were done with Cytobank and FlowJo. Antibodies with positive signals were validated by FACS as summarized in (Table S1). We applied the cluster analysis programs SPADE and viSNE used for CyTOF51 to create a differentiation panel of the resultant screened markers.

### RNA extraction, purification, and transcriptome sequencing

RNA isolation of sorted cells is based on a previous work^52^: 20,000 cells of each sequenced population were sorted in 750 μl Trizol-LS (Invitrogen #10296010) using a 100 μm nozzle. Cells were vortexed and incubated for 10 minutes at room temperature prior to freezing in -80 C. After thawing, cells were washed with chloroform, and RNA was isolated according to the manufacturer’s directions, with minor modifications. Linear polyacrylamide (LPA) carrier (Sigma Aldrich # 56575-1ML) was added to enhance recovery of RNA, followed by DNase I treatment per manufacturer’s instructions (Qiagen RNeasy micro kit, # 74004).

Tissue sections: Tissue dissections were stored at -80 C, resuspended in Trizol and liquid Nitrogen to minimize RNA degradation, and then homogenized into a soft powder using vigorous mechanical disruption with a mortar and pestle. Homogenized-frozen tissue was allowed to thaw. RNA was isolated according to the manufacturer’s directions, with minor modifications. Linear polyacrylamide (LPA) carrier was added to enhance recovery of RNA, followed by DNase I treatment per manufacturer’s instructions (Promega RQ1 DNase).

Recovered RNA was then analyzed by an Agilent 2100 Bioanalyzer for quality analysis prior to library preparation. cDNA libraries were then prepared from high quality samples (RIN > 8) using Ovation RNA-seq v2(Nugen); NEBnext DNA Master Mix for Illumina (New England Biolabs) and standard Illumina adapters and primers from IDT. Barcoded library samples were then sequenced on an Illumina NextSeq 500 (2x150bp, producing an average of 15 million reads/cell population). The 23 endpoint populations were sequenced, in addition to CP2 which was composed of all live cells. CP17 was composed of dead cells and cell debris, and therefore was excluded in the population analysis.

### Cytotoxicity and phagocytosis assays

For in vitro cytotoxicity assays we used a FACS-based cytotoxicity assay^19,53^. Isolated *B. schlosseri* cells were labeled using CFSE (5uM; Life technologies) and Far Red dye (1uM; Life technologies) for 30 minutes at 18° C to distinguish effector and target cells and washed twice in staining media: 3.3x PBS, 2% FCS and 10 mM Hepes. Cells were incubated overnight in 96 well U shaped plates at 18 degrees at different effector: target ratios in staining media. After adding Propidium Iodide (PI) to wells for cell viability separation, wells were analyzed by FACS. Spontaneous lysis was measured as the percentage of PI positive cells in gated target cells in wells without effector cells, and sample lysis was quantified as the percentage of PI positive cells in gated target cells. Specific lysis was calculated as follows: Specific lysis %=100x((sample lysis-spontaneous lysis)/(100-spontaneous lysis)). The gating on FACS analysis was on two dimensional plots due to the natural fluorescence of *B. schlosseri* cells. For BHF blocking assays, anti-BHF polyclonal mouse serum was used 1:200, or mock serum as control. Colonies with known fusion/rejection outcomes from our mariculture facility were used to measure allogeneic cytotoxicity and phagocytosis. Cytotoxicity and phagocytosis assays were done in triplicates and the results represent one experiment of three performed.

For *in-vitro* allogeneic phagocytosis assays we also developed a FACS-based phagocytosis assay. The labeling and incubation of cells were done as for cytotoxicity assay. Phagocytosis was measured as the double positive cells in the FACS plots of the two labeling markers. Due to the natural fluorescence of *B. schlosseri* cells, the level of double positive cells in wells with separation of each one of the labels was reduced from the double positive in the experimental wells (about 5% background). For validation of phagocytosis, cells were sorted and observed by confocal microscopy. To compare cytotoxicity to phagocytosis, the optimal ratio (1:1) for allogeneic phagocytosis was prepared and compared to cytotoxicity at the same wells, using 10^5^ cells of each labeled group per well.

Bacteria and beads phagocytosis assays: cells (10^5^ cells/200 μl) were incubated overnight at 18°C in a 2:1 ratio of beads: cells using Fluoresbrite^®^ YG Carboxylate Microspheres 1.00μm (Polysciences). For bacterial phagocytosis, the marine bacteria *Vibrio diazotrophicus* was heat inactivated in 95° C for 5 minutes, and labeled with Alexa Fluor 647 (Invitrogen #A20006). For bacteria phagocytosis, cells (10^5^ cells/200 μl) in a 2:1 bacteria:cells ratio, were incubated overnight at 18° C. The analysis of cells positive to beads or cells positive to bacteria was done by flow cytometry in two dimensions - on the green channel for beads or far-red channel for bacteria (Fig. 4A). Analysis of flow cytometry data was accomplished using FlowJo V10 (FlowJo).

### Cell Transplantation

Cell were sorted directly into staining media (3.3x PBS based; 75% of final volume) in order to minimize cellular stress using a 100 μm nozzle. Cells were then labeled with DiD or DiO membrane dye in order to visualize or identify the transplanted population, then suspended in a final concentration of 10^5^ cells/μl. 10^5^ cells were transferred into recipient colonies via zooid injections near the heart using a manual microinjector and micromanipulator (Narishige, Japan). After the transplantation experiment, the analysis for the pigment cell chimerism and differentiation assays (Fig. 2b-c) was done in a single blind manner 20 days following transplantation. The experiment represents 1 of two performed. Experiment-1 N=16, Experiment-2 N=42 (15-control population, 16-cHSC, 11-uninjected). On day 21 of Experiment-2 the changes in the pigment cells were validated by FACS due to the lower auto-fluorescence on the yellow channel of blue pigment cells compared to the orange pigment cells (Fig. S1). For general cell differentiation analysis, transplanted colonies were pooled for the analysis by FACS into two pools from each group, having at least 5 transplanted animals in each pool (Fig. 2d-e). For localization assays two sets of experiments were performed: a total of 14 colonies labeled with CFSE on ibidi 50 mm, uncoated μ-dishes were transplanted with DiD labeled cells (Fig. 2f, S6a).

### Electron Microscopy

Colonies were fixed overnight in 1.5% glutaraldehyde buffered with 0.2 M sodium cacodylate, 1.6% NaCl, pH 7.4. After washing in buffer and post-fixation in 1% OsO4 in 0.2 M cacodylate buffer plus 1.6% NaCl, specimens were dehydrated and embedded in Araldite. Sections were counterstained with toluidine blue, and for EM sections were stained for contrast with uranyl acetate and lead citrate. Micrographs were taken with a FEI Tecnai G2 electron microscope (operating at 100 kV).

### Differential expression analysis

Determination of gene counts was performed using a Snakemake^54^ pipeline. An outline of the steps is as follows: i) low quality bases and adapter sequences were removed using Trimmomatic^55^ (version 0.32) ii) overlapping paired end reads were merged using FLASH^56^ (version 1.2.11) iii) reads were aligned to the UniVec Core database using bowtie2^57^ (version 2.2.4) to remove biological vector and control sequences, iv) reads were aligned to the *Botryllus schlosseri* transcriptome with BWA^58^ (“mem” algorithm, version 0.7.12), v) aligned reads were sorted and indexed using samtools, PCR duplicates removed using PICARD (“MarkDuplicates” tool, version 1.128) and then transcript level counts directly counted from the BAM file.

Differential expression was performed using edgeR^59^. In detail: the gene counts were compiled into a tabular format and loaded into R. Genes were retained with at least five counts per million in at least 80% of the smaller number of the compared samples. A simple model was used to compare the two sets of populations, with p-values adjusted using the Benjamini-Hochberg method to produce a false discovery rate (FDR). FDRs less than 0.05 were called as being differentially expressed. The comparisons between cell populations were performed in a one-vs-all approach, followed by selective aggregating of similar populations. Initially each population was compared to all others (except for CP2 which was an aggregate of all cells). Then all sets of two populations were compared to the remaining, with the best two being grouped together. The metric used is: if A is the set of genes found to be differentially expressed in population A vs all others, and n(A) is the number of genes in that set, then find the maximum of n(AB) - n(A or B). This attempts to find those populations that when grouped together are more distinct from all others compared to the populations individually. After this grouping was performed multiple times, the maximum was no longer above zero, and a new metric was used. This method was used to find the two groups of populations that maximized n(AB)^2^/(n(A or B) + 1). This aimed to find the populations that fractionally were the most similar. The mergings are as follows: CP33+CP34, CP3+CP23, CP8+CP20, CP21+CP22, CP24+CP26, CP25+CP33+CP34, CP12+CP14, CP5+CP32, CP29+CP36, CP19+CP31, CP15+CP16, CP30+CP35. The number of characteristic, highly expressed genes found per population(s) relative to the other populations varied from 0-2229 (mean 232), with only 23% possessing homology to mouse or human genes with the remaining 77% of these genes being mostly *B. schlosseri* or tunicate specific.

For visualization, the 250 genes with the largest weights in the first 11 principal components (explaining 90% of the variance in mean-adjusted log-transformed gene counts) were used to cluster the different cell populations in a heatmap (Fig. S4a). Many of the populations have visually similar gene expression patterns (for example: CP3, 5 and 23; and CP25, 33 and 34), and in general, samples that were adjacent in FACS-space are also similar in expression-space (Fig. S4b, 67% of neighbors in agreement) indicating a correlation between gene expression profile and morphology and marker expression.

### Geneset Activity Analysis on Gene Expression Commons

Gene Expression Commons (https://gexc.riken.jp) provides gene expression dynamic-range and threshold to distinguish active expression from inactive expression for each gene by computing a massive amount of publicly available microarray data24. Once a user submits raw microarray data of a particular cell type, the Gene Expression Commons returns gene expression activity referred to the dynamic-range for each gene, and if expression level exceeds the threshold, the gene is labeled “active gene”. In the Weissman lab, we purified and generated microarray data for each step of adult mouse hematopoiesis as well as several stromal cell types. A gene expression profile of each cell type becomes public domain (or available to the public) through the Gene Expression Commons platform.

For this project, we developed a new function on the Gene Expression Commons platform entitled Geneset Activity Analysis. A user can create a Geneset, a set of genes of interest. Then the significance of the association between a given gene set and a set of active genes in the cell type is examined by Fisher’s exact test.

## Supplementary Information

**Supplementary Figure 1.**
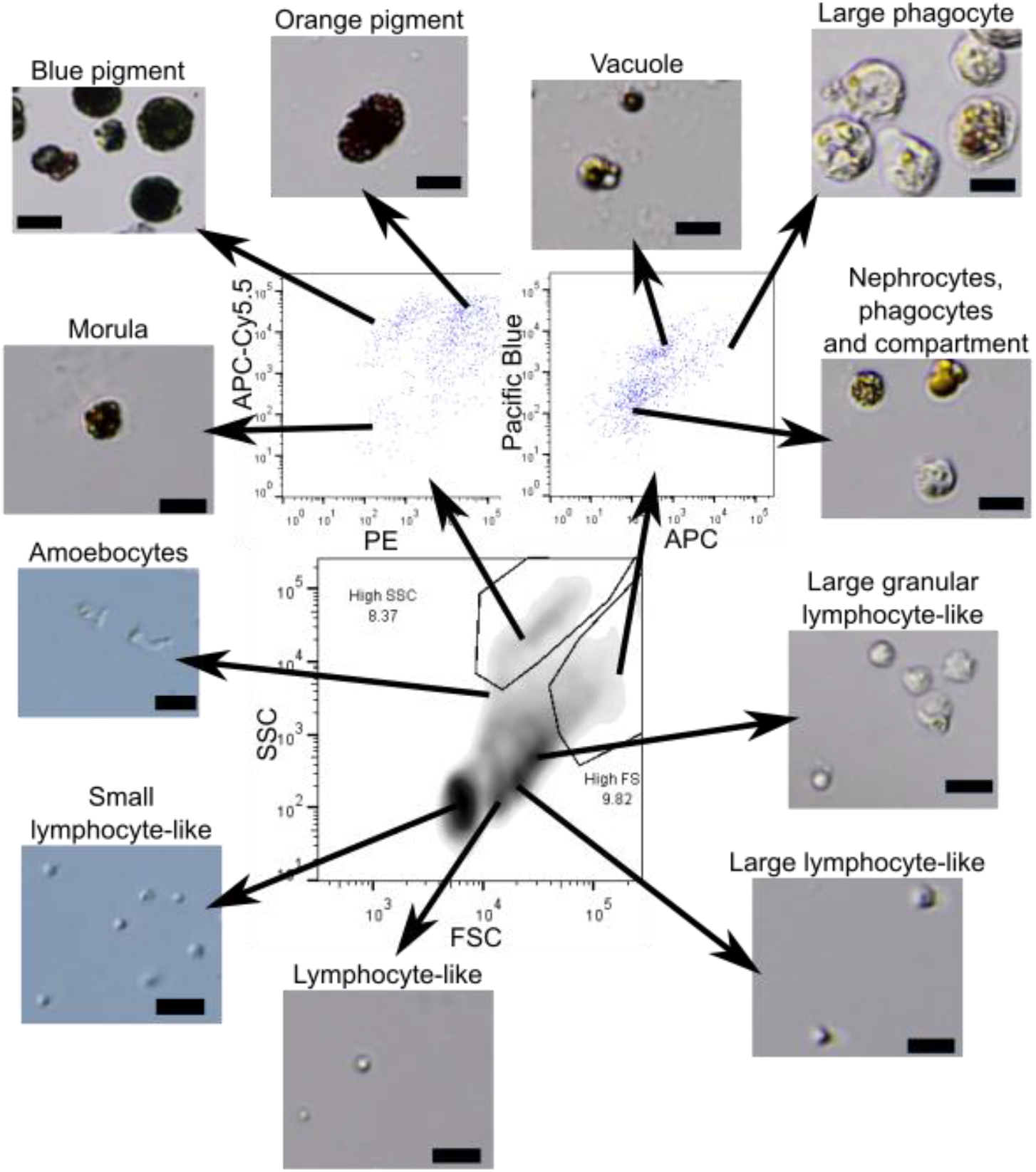
Morphologically sorted cell types and names. Sorting based on FCS/SSC in the lower panel, and natural fluorescence in the upper panels. The analysis is after gating PI negative cells (live cells). The specific excitation laser and optical filter for emission measurements was as follows [excitation laser in nm and filters stated as long pass (LP) and band pass (BP)]: 488nm for FSC and SSC, 488nm(550LP 575/25BP)-PE, 405nm(450/50BP)-Pacific Blue, 633nm(660/20BP)-APC, 633nm(690LP 710/50BP)-APC-Cy5.5. Nomenclature based on ^14,28^. Scale bar 20 μm.

**Supplementary Figure 2.**
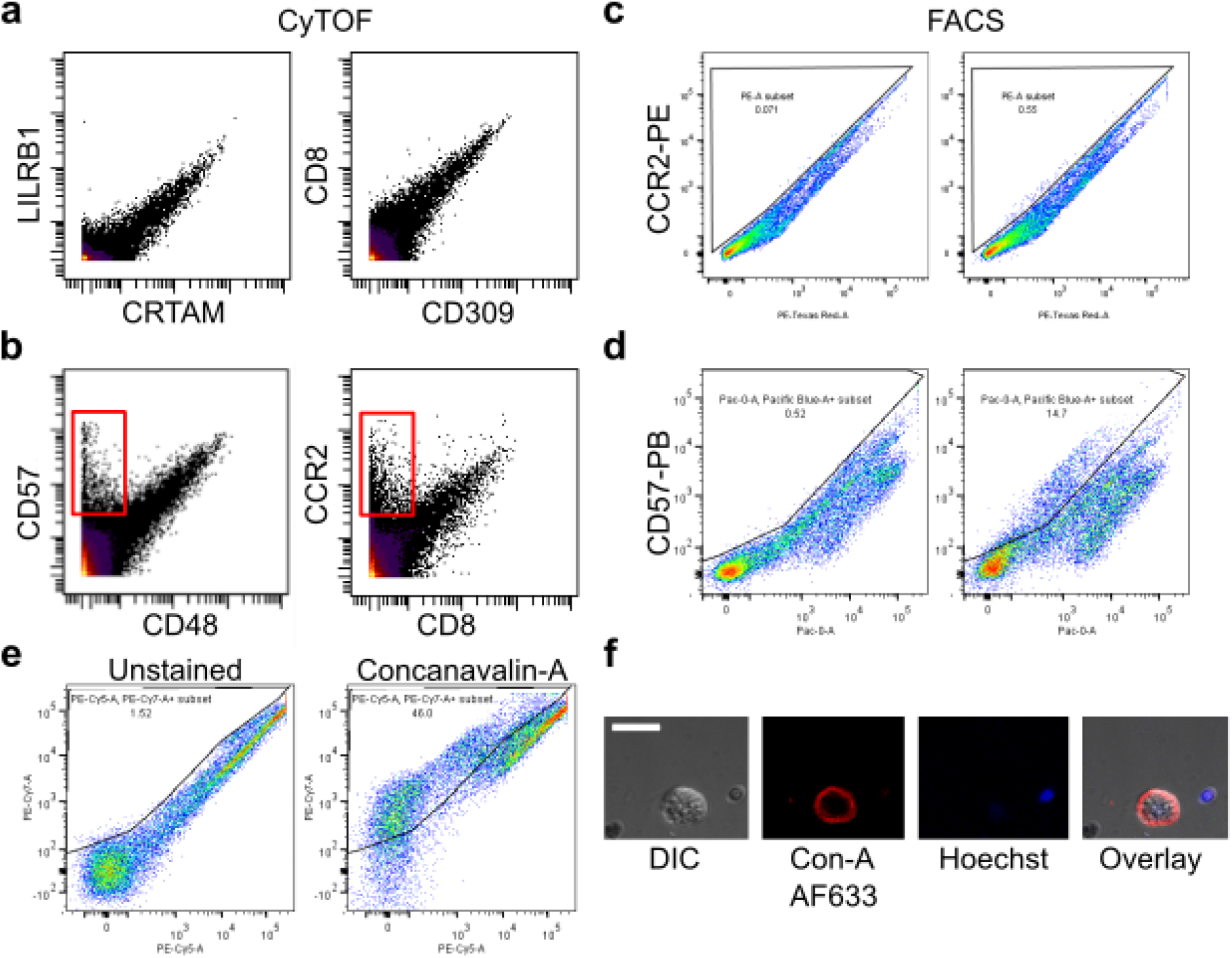
Screen for differentiation markers of cell populations for FACS based sorting. **a-b,** Examples of screened antibodies by CyTOF Mass cytometry with *B. schlosseri* cells analyzed by two dimensional mass spectrometry. **a,** Examples of antibodies considered as nonspecific binders due to the same binding patterns by different antibodies. **b,** Examples of antibodies considered as specific binders due to the cell population that is bound by one antibody but not the other (labeled in red). **c-d,** Examples of validation screen by flow cytometry of antibodies with specific binding by CyTOF, done in two dimensional fluorescence excited by the same laser due to autofluorescence. **c,** Negative validation of CCR2 labeled with PE. **d,** Positive validation of CD57 labeled with Pacific Blue. **e,** Example of positive differential labeling by the lectin Concanavalin-A in PE-Cy7 by FACS. **f,** Confocal imagery of membrane Concanavalin-A labeling in Alexa Fluor-633 in red and Hoechst DNA labeling in blue. Scale bar 20 μm.

**Supplementary Figure 3.**
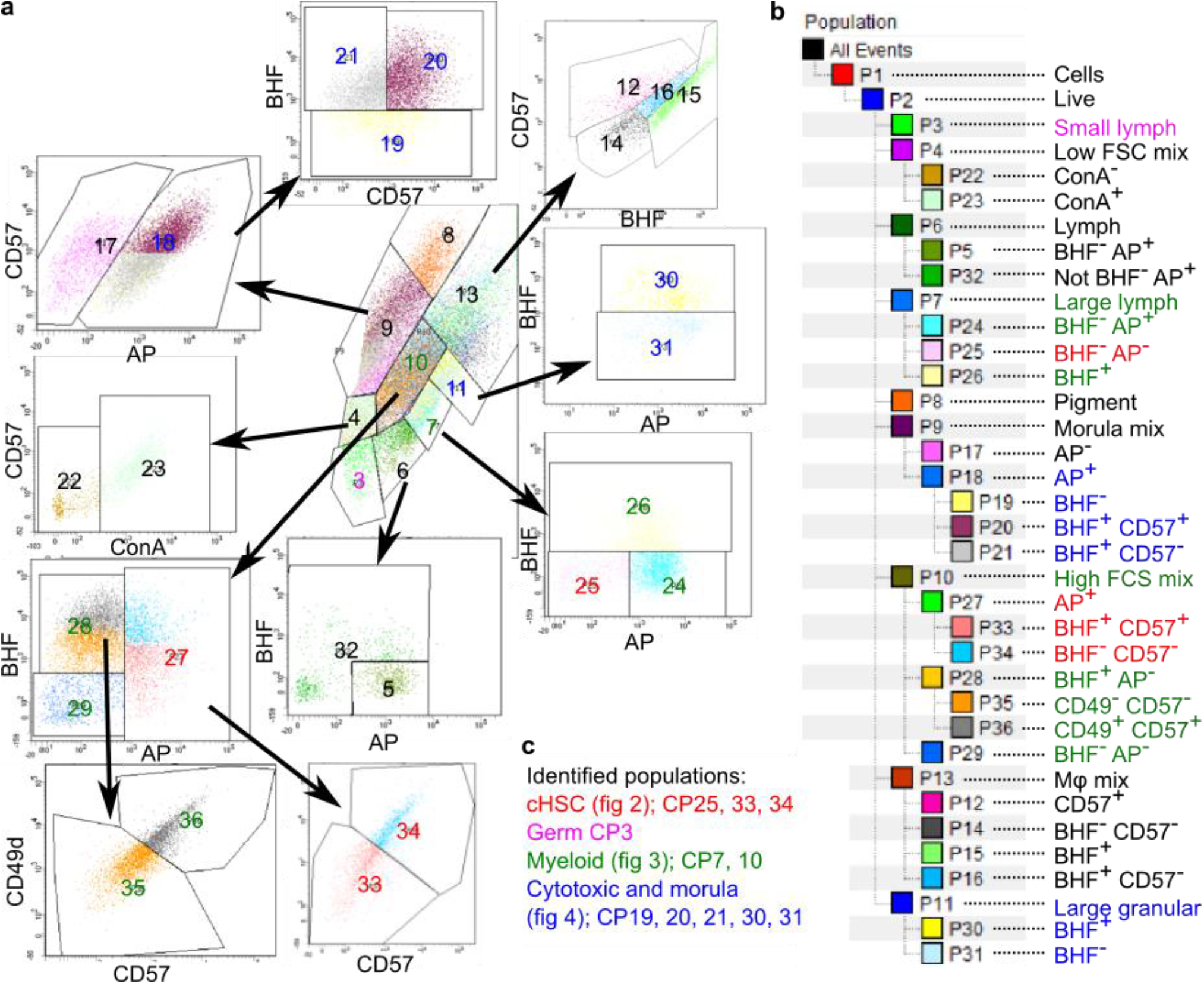
Thirty Four cell population panel. **a,** Sorting panels of 34 cell populations using: FSC, SSC, CD49d, CD57, Concanavalin-A (ConA), BHF and AP, after gating PI negative cells (live cells). Central panel is FCS/SSC from which additional populations were differentiated. The specific excitation laser and optical filter for emission measurements was as follows [excitation laser in nm and filters stated as long pass (LP) and band pass (BP)]: 488nm(505LP 530/30BP)-AP, 488nm(755LP 780/60BP)-CD49d, 405nm(450/50BP)-CD57, 633nm(660/20BP)-ConA, 633nm(755LP 780/60BP)-BHF. **b,** Hierarchy of sorted cell populations by main parameters of differentiation. **c,** Color key (reflected in subparts **a-c**) for different populations identified in this study and the Figures that further describe them.

**Supplementary Figure 4.**
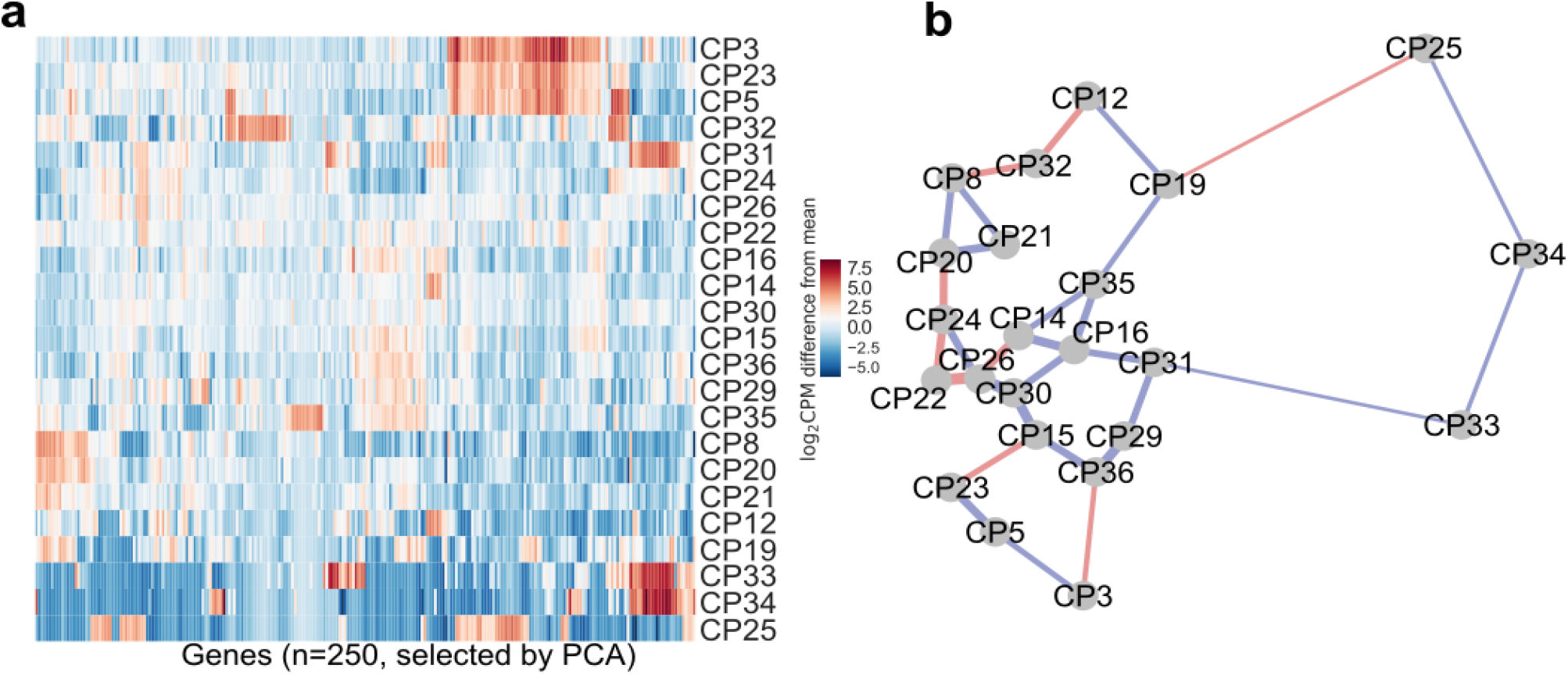
*B. schlosseri* cell population clustering based on transcriptome analysis. **a,** 250 genes with the largest weights in the first 11 principal components (explaining 90% of the variance in mean-adjusted log-transformed gene counts) were used to cluster the different cell populations in a heatmap. **b,** Transcriptome sequencing of *B. schlosseri* cell populations compared to FACS analysis. 2D projection of the distances between transcriptomes of cell populations based on all differential genes. Lines are drawn between the nearest two neighbors. Blue: corresponds to the FACS adjacency of the populations in the differentiation panel, in red: genetic level proximity that is not predicted by FACS panel adjacency. 20/30 genetic level cellular populations’ proximity were predicted by FACS. Width of lines is inversely proportional to the distances.

**Supplementary Figure 5.**
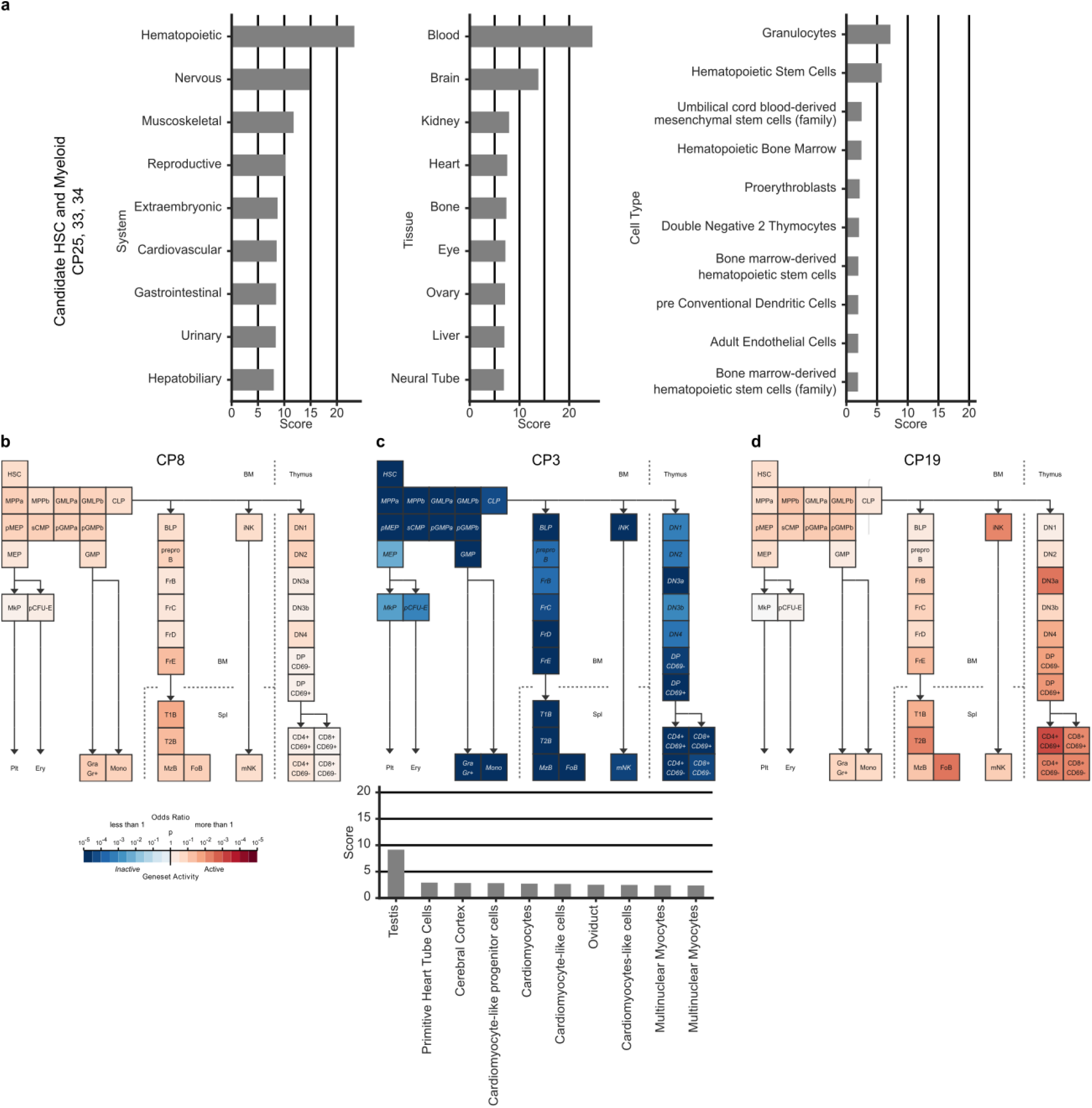
Gene expression of *B. schlosseri* cell populations. **a,** Enrichment scores of the top ten systems (left), tissues (center), and cell types (right) of annotated genes up-regulated in CP33, 34 and 25 using the GeneAnalytics tool compared to human. In systems the hematopoietic system is the highest score, in tissues the blood is the highest score, and within the cells, the granulocytes and HSC have the highest scores. **b-d,** Geneset Activity Analyses using the Gene Expression Commons tool on a mouse hematopoiesis model of different *B. schlosseri* cell populations. **b,** Analysis of CP8 (pigment cells) based on 12 significantly upregulated genes, showing that CP8 is part of the hematopoietic system with gene activity resembling known cell type. **c,** Analysis of CP3 (small cells) based on 235 significantly upregulated genes, showing that this population is significantly not part of the hematopoietic system. On the bottom: enrichment scores of the top ten tissues using CP3 genes by GeneAnalytics tool compared to human, the highest score is the testis suggesting this population is gonadal population. **d,** Analysis of CP19 (enriched with morula cells) based on 96 upregulated genes (p<0.25), showing that CP19 has gene activity as cells in the lymphoid lineage using Geneset analysis.

**Supplementary Figure 6.**
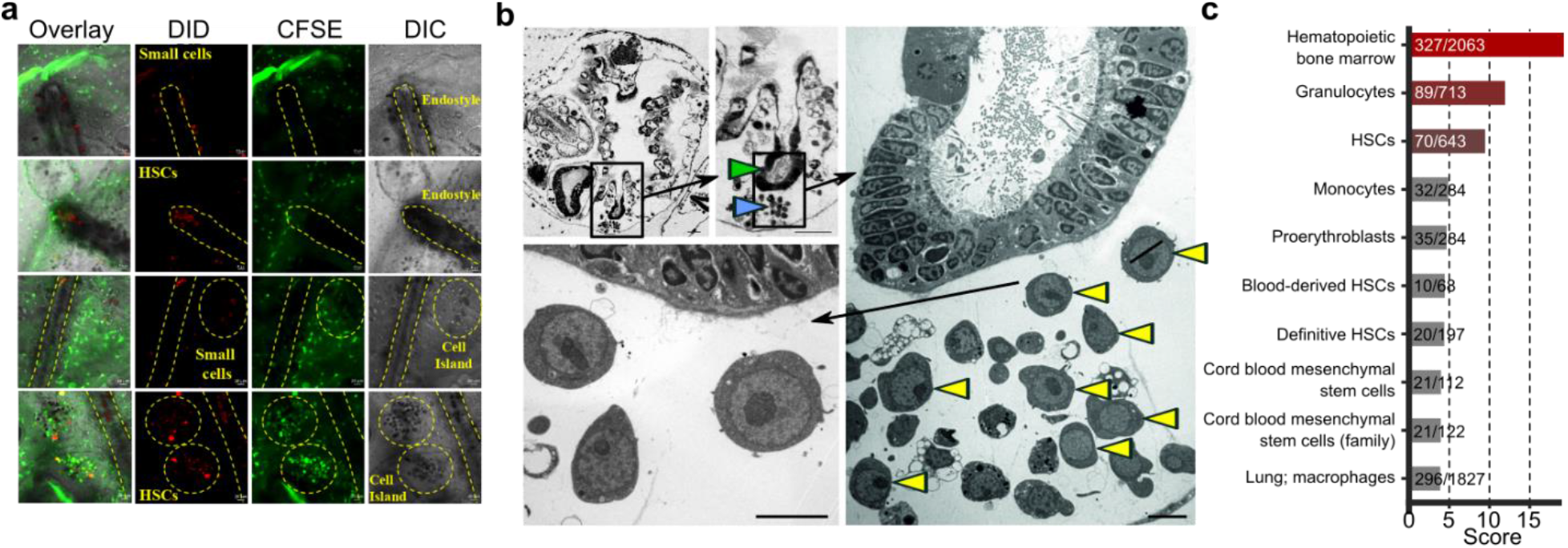
Sub-endostylar sinuses as an HSC niche. **a,** Candidate HSC (cHSC) population and a control population (CP3) were isolated, labeled with DiD and transplanted into CFSE labeled compatible colonies, *in-vivo* tracing of transplanted cells migration was used to identify niches. Five to ten days afterwards, only the cHSC populations migrated and aggregated in the sub-endostylar sinuses (endo-niche). Both of the populations aggregated at the cell Islands. **b,** Transverse sections of an adult zooid counterstained with toluidine blue (top two left) where endostyle (green arrow) and endo-niche (blue arrow) are enlarged (Scale bar: 30 μm). Electron microscopy section of the same animal endostyle and endo-niche (right and bottom enlarged). Red arrows indicate cells with hemoblast (HSC) morphology that are enriched within the endo-niche (Scale bar: 5 μm). **c,** Enrichment scores of the top ten tissues/cell types and number of genes matching (compared with total in that group) based on the blood-associated differentially expressed endostyle genes.

**Supplementary Figure 7.**
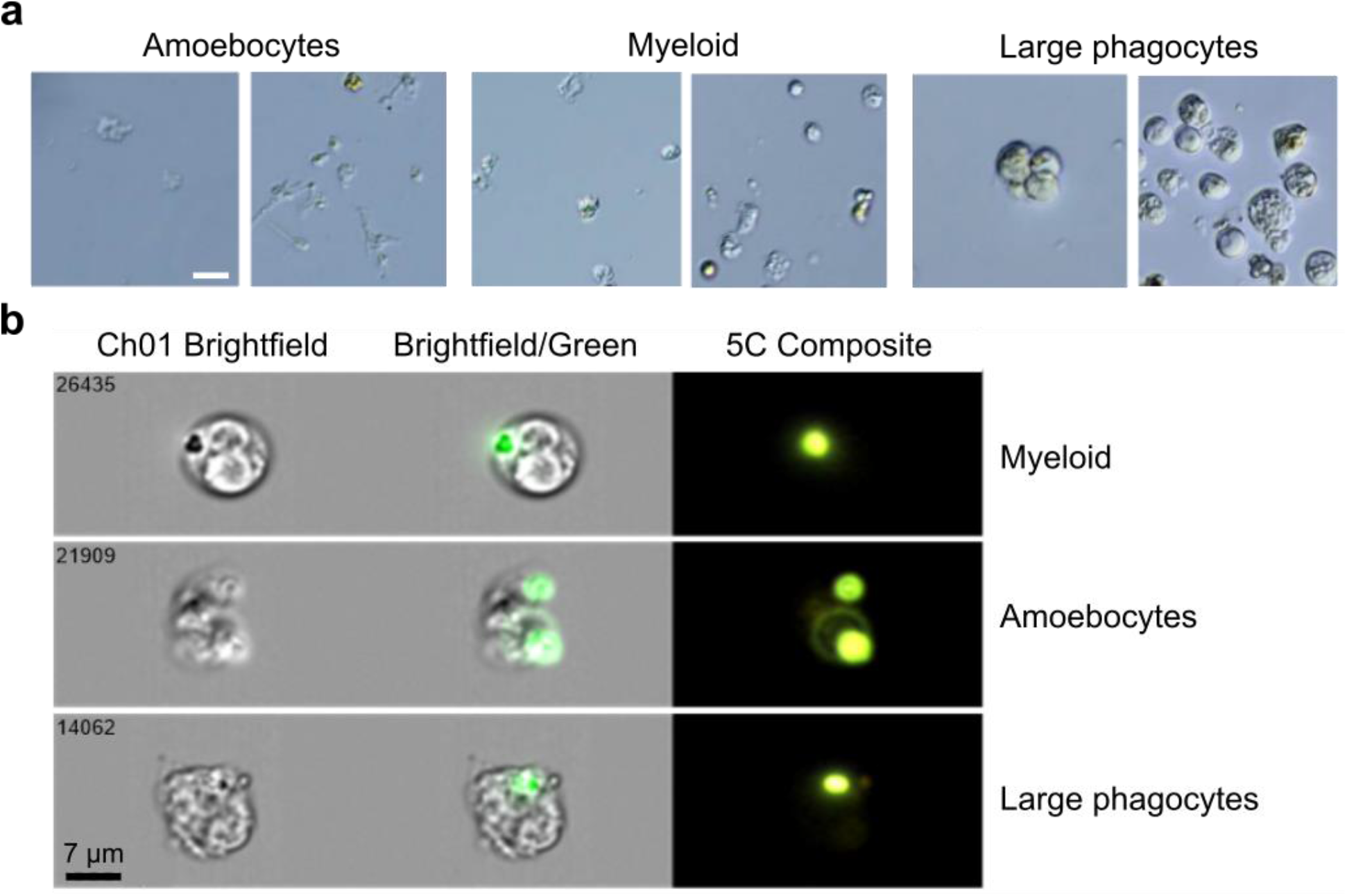
*B. schlosseri* Myeloid Lineage Phagocytic Population. **a,** Live images of the three isolated phagocytic populations, scale bar 20 μm. **b,** Phagocytosis of green fluorescent beads by *B. schlosseri* cells analyzed by ImageStream. The positive cells have mainly morphology of: amoebocytes, myeloid cells, and large phagocytes. Representative images of the three phagocytic populations after engulfment of beads. Scale bar 7 μm.

**Supplementary Figure 8.**
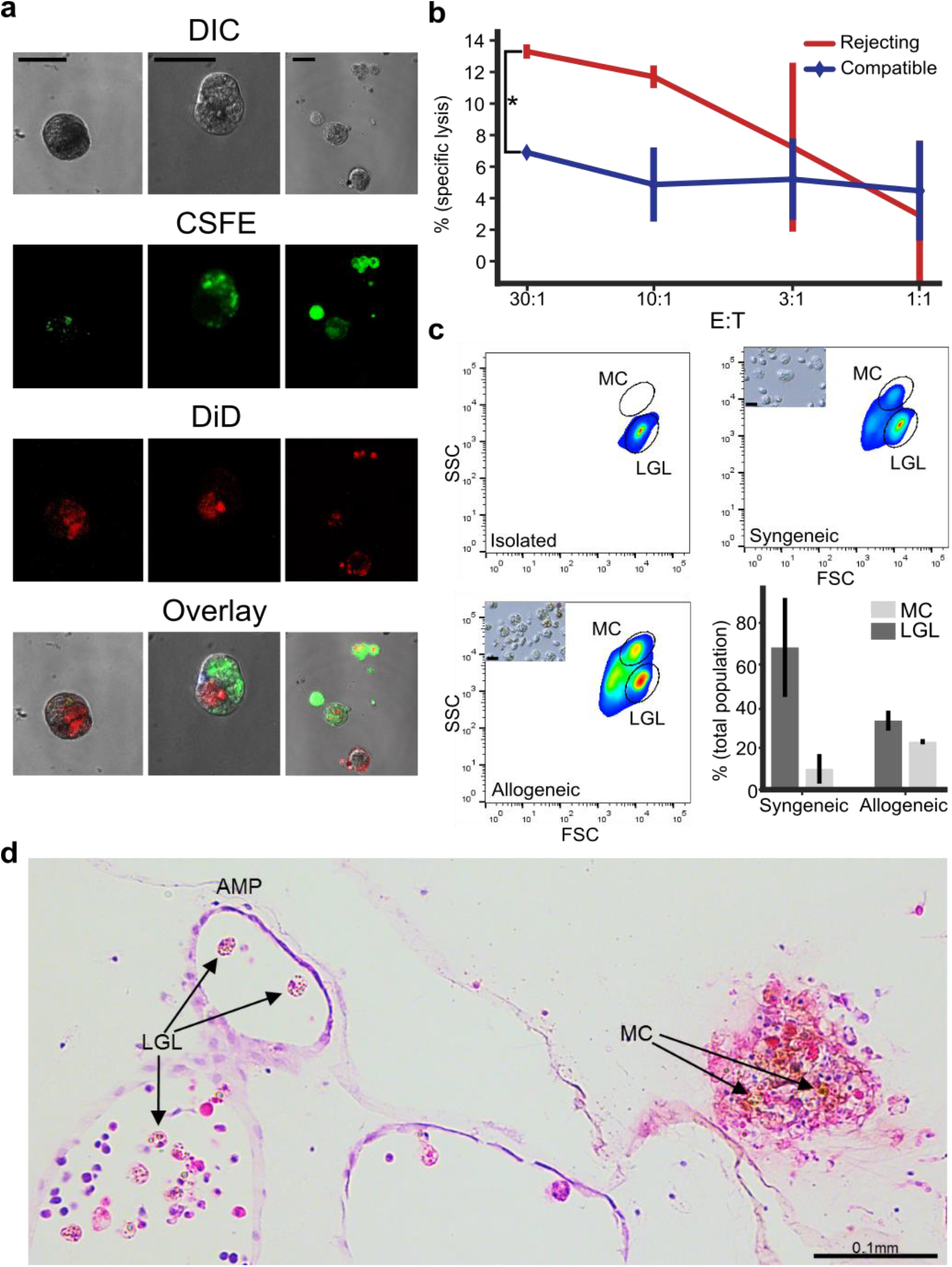
Cytotoxicity and the two morphs of morula cells (MC) at point of rejection (POR). **a,** Confocal imagery of phagocytosis assays to validate the allogeneic engulfment. In the panels colony labeled with CFSE in green and colony labeled with DiD in red after allogeneic phagocytosis assay. Large phagocytic cells can be seen after engulfment of allogeneic cells or vesicles. Scale bar 20 μm, **b,** Example of cytotoxicity assay in different Effector to Target ratios (E:T), where the targets are compatible or rejecting colony cells to the effector colony. In the rejecting colony, specific lysis is significantly higher. **c,** LGL cells were isolated (upper left) and incubated overnight either in syngeneic (upper right) or in allogeneic challenge (lower left). FSC/ SSC analysis of LGL cell (lower population) and MC (upper population). Small caption of light microscopy pictures of cells after the incubation. (Lower right) Analysis of LGL and morula cells in syngeneic compare to allogeneic challenge. Bars= Standard Deviation. **d,** An H&E section of *B. schlosseri* colonies undergo rejection. In the ampule (AMP) the inactivated form of cytotoxic MC/ large granular lymphocytes like cells (LGL) can be observed (left). On the other hand the activated form with the brown pigmentation of MC can be observed at POR (right).

**Supplementary Table 1.** Screened antibodies by CyTOF for binding of *B. schlosseri* cells. Symbol: represents the element, mass: is the elemental isotope mass, and antigen: is the human antigen against which the antibody was produced. CyTOF column represents whether the *B. schlosseri* cell population was positive or not, low means less than 1% of cells were positive. FACS column represents whether there was binding by the same antibody clone by flow cytometry.

| Symbol | Mass | Panel1 antigen | CyTOF | FACS | Panel2 antigen | CyTOF | FACS |
| --- | --- | --- | --- | --- | --- | --- | --- |
| Pd/Cd | **110-114** | CD3 | Yes | No |  |  |  |
| In | **113** | CD7 | Yes | No |  |  |  |
| In | **115** | CD45 | Yes | No |  |  |  |
| I | **127** |  |  |  |  |  |  |
| La | **139** | CD57 | Yes (low) | Yes (~14%) | CD2 | Yes (low) | No |
| Pr | **141** | NKp46 | No |  | CD61 | No |  |
| Nd | **142** | CD48 | No |  |  |  |  |
| Nd | **143** |  |  |  |  |  |  |
| Nd | **144** | CCR5 | Yes | N/A | CD94 | Yes (low) | N/A |
| Nd | **145** |  |  |  | LILRB1 | No |  |
| Nd | **146** |  |  |  | CD309 | No |  |
| Sm | **147** |  |  |  | CD8 | No |  |
| Nd | **148** | KIR3DL1 | No |  | CRTAM | No |  |
| Sm | **149** | DNAM1 | No |  |  |  |  |
| Nd | **150** |  |  |  |  |  |  |
| Eu | **151** | NKG2D | No |  | CD84 | No |  |
| Sm | **152** | TNFR2 | No |  |  |  |  |
| Eu | **153** | NKG2C | Yes (low) | N/A |  |  |  |
| Sm | **154** | NKp44 | No |  | Notch1 | No |  |
| Gd | **155** | CRACC | No |  |  |  |  |
| Gd | **156** | KIR2DL3 | No |  |  |  |  |
| Gd | **157** |  |  |  |  |  |  |
| Gd | **158** | CD161 | No |  | clec12A | No |  |
| Tb | **159** |  |  |  | CD11c | No |  |
| Gd | **160** | NKp30 | No |  | 2B4 | No |  |
| Dy | **161** | CD15 | No |  | CD4 | Yes | No |
| Dy | **162** | CD49d | Yes | Yes (~28%) |  |  |  |
| Dy | **163** | CD16 | No |  | KIR2DLS1 | No |  |
| Dy | **164** |  |  |  |  |  |  |
| Ho | **165** |  |  |  |  |  |  |
| Er | **166** | TIGIT | No |  | KIR3DL2 | No |  |
| Er | **167** | CD27 | No |  | CCR7 | No |  |
| Er | **168** | KIR2DL1 | No |  | CCR2 | Yes | No |
| Tm | **169** |  |  |  |  |  |  |
| Er | **170** | CD11a | No |  | CD11b_act | No |  |
| Yb | **171** |  |  |  |  |  |  |
| Yb | **172** | KIR2DL4 | No |  | CD34 | Yes (low) | N/A |
| Yb | **173** |  |  |  | CD33 | No |  |
| Yb | **174** | CD144 | No |  | NKG2A | No |  |
| Lu | **175** | KLRG1 | No |  |  |  |  |
| Yb | **176** |  |  |  | CD56 | Yes (low) | N/A |
| Ir | **191/193** | DNA | All cells |  | DNA | All cells |  |
| Pt | **195** | cisplatin | No |  |  |  |  |

**Supplementary Table 2.** Flow cytometry binding of *B. schlosseri* cells. The column of % positive cells is the percentage of the cells that were positively labeled by the marker. For Alkaline Phosphatase (AP) the high represent cells that labeled strongly and mid-cells that are positively labeled but not high.

| Lectin | Fluorophore | % positive cells |
| --- | --- | --- |
| ConA | AF-633 | 45% |
| PNA | PE-Cy7 | 16% |
| UEA | PE-Cy7 | 30% |
| **Additional markers** |  |  |
| Alkaline Phosphatase | Green | 9% high 25% mid |
| Anti-BHF polyclonal | AF-647 | 35% |

## Additional supplementary files

**Supplementary movie 1: Budding cycle and take-over.** Three day time lapse acquisition of buds’ development in two *B. schlosseri* colonies (colony 1 and colony 2). At time lapse’s 35th hour point, the zooids of colony 2 are getting absorbed and replaced by their buds. Taken on a Kyence BZ-X700 every 18 minutes during a 60 hour period.

**Supplementary movie 2: Vascular fusion between two compatible *B. schlosseri* colonies.** Time-lapse acquisition of vascular fusion between two *B. schlosseri* colonies (colony 1 and colony 2). At the time lapse’s 3 hour and 10 minute mark the ampullae in the center touch, than at the 6 hour 40 minute mark they fuse. Movies of blood flowing from one colony to the other are shown after. Taken on a Kyence BZ-X700 every 10 minutes during an 18 hour period.

**Supplementary movie 3: Differential labeling of colonies.** Live colonies labeled on ibidi 35mm micro dishes with either CFSE (green) or cell tracer far red (red). Also an example of fused colonies. Taken by confocal microscopy.

**Supplementary table 3: Gene expression by populations.** Reads per kilobase of transcript per million for each gene of each isolated endpoint population (fig. 1e-g, S3,S4). On the right side is the results of the differential expression analysis for each populations or cluster of populations with adjusted (Benjamini-Hochberg) p-values <0.05 and <0.25. Differentially up-regulated=1, differentially down-regulated=-1, no differential expression=0.

**Supplementary table 4: Genes sets used throughout the study.** List of the genes used in the analysis in Gene Commons Tool and the Gene Analytics. If gene name appears more than once it means that there are more than one *B. schlosseri* gene model that is associated with this gene and was upregulated.

